# Analysis of Genetically Determined Gene Expression Suggests Role of Inflammatory Processes in Exfoliation Syndrome

**DOI:** 10.1101/2020.12.17.423318

**Authors:** Jibril B. Hirbo, Francesca Pasutto, Eric R. Gamazon, Patrick Evans, Priyanka Pawar, Daniel Berner, Julia Sealock, Ran Tao, Peter S. Straub, Anuar I. Konkashbaev, Max Breyer, Ursula Schlötzer-Schrehardt, André Reis, Milam A. Brantley, Chiea C. Khor, Karen M. Joos, Nancy J. Cox

## Abstract

Exfoliation syndrome (XFS) is an age-related systemic disorder characterized by excessive production and progressive accumulation of abnormal extracellular material, with pathognomonic ocular manifestations. It is the most common cause of secondary glaucoma, resulting in widespread global blindness. We performed Transcriptomic Wide Association Studies (TWAS) using PrediXcan models trained in 48 GTEx tissues to identify genetically-determined gene expression changes associated with XFS risk, leveraging on results from a global GWAS that included 123,457 individuals from 24 countries. We observed twenty-eight genes in a three-Megabase chr15q22-25 region that showed statistically significant associations, which were further whittled down to ten genes after additional statistical validations. In experimental analysist of these ten genes, mRNA transcript levels for *ARID3B, CD276, LOXL1, NEO1, SCAMP2,* and *UBL7* were significantly decreased in iris tissues from XFS patients compared to control samples. Genes with genetically determined expression changes in XFS were significantly enriched for genes associated with inflammatory conditions. We further explored the health consequences of high susceptibility to XFS using a large electronic health record and observed a higher incidence of XFS comorbidity with inflammatory and connective tissue diseases. Our results implicate a role for connective tissues and inflammation in the etiology of XFS. Targeting the inflammatory pathway may be a potential therapeutic option to reduce progression in XFS.

## Introduction

Exfoliation syndrome (XFS) is an age-related systemic disorder characterized by excessive production and progressive accumulation of abnormal extracellular material, with pathognomonic ocular manifestations.^1,2^ It is the most common cause of secondary glaucoma, resulting in widespread global blindness.^3^ In addition to ocular manifestations, exfoliation syndrome deposits have been observed in visceral organs, such as the lung, kidney, liver and gallbladder.^2,4^ In addition to elastic tissue disorders, XFS has also been associated with increased risk of vascular diseases.^5–7^ Associations of XFS to several systemic biomarkers of inflammation, including complement components and homocysteine, have also been reported.^3,8,9^

Genetic mechanisms have substantial influence on XFS etiology as evidenced in family and twin studies.^10,11^ There have been eight genome-wide association studies (GWAS) of XFS,^7,12–18^ three of which include meta-analysis,^7,12,13^ that have cumulatively identified >60 associated genetic variants. The largest meta-analysis of XFS involved >123,000 individuals (13,620 XFS cases, 109,837 controls) from 24 countries across six continents and identified seven loci with the strongest association signal in chromosome 15 near the lysyl oxidase-like 1 gene (*LOXL1*). The signal on chr15 involved 54 potential causal variants. Overall, (i) two missense variants in *LOXL1,* rs1048661 (encoding *LOXL1* p.Leu141Arg) and rs3825942 (p.Gly153Asp), are likely to confer risk of developing XFS, with very high heterogeneity across populations because the alleles show an effect reversal,^12,13,16,19–22^ (ii) the associated variants in the locus showed population-specific frequency and linkage disequilibrium (LD) patterns,^12,16,21^ (iii) haplotypes that carry the risk alleles depending on the population are correlated with reduced *LOXL1* expression levels, however, (iv) no clear functional effects for the haplotypes that represent the two variants have been shown.^7,13,23,24^ The non-coding variants associated with XFS at this chr15 locus could confer regulatory effects. Some of these non-coding variants regulate expressions of the sentinel *LOXL1* and the neighboring *STRA6* gene.^7,25,26^

After considering all the reports on genetic architecture of XFS to date, we hypothesize that analysis of the contribution of the genetically-determined component of gene expression to XFS risk can provide a powerful method to elucidate genes involved in XFS. We used a genebased TWAS method, PrediXcan^27^, implemented on GWAS summary statistics (Summary PrediXcan; S-PrediXcan)^28^ to identify genetically determined gene expression traits associated with disease risk. Models were trained on 48 GTEx tissues to estimate the correlation between genetically-determined gene expression and XFS risk, leveraging on XFS GWAS summary statistics from a previously reported multi-ethnic study.^13^ The phenomenon of TWAS association with multiple signals within the same locus can be a statistical artifact of the correlation due to LD between SNPs that are separately predictive of the measured expression of physically colocolized genes^32^ hampering the ability to prioritize the true causal gene(s). To address this limitation, we performed sequential conditional analysis in each tissue, starting with the gene that was the strongest signal in the initial PrediXcan analysis. In addition, we sequentially rebuilt prediction models excluding variants in models of other genes in the loci that were in LD with any variants of the strongest signal. We also analyzed individual-level GWAS data from four additional European ancestry populations, two German, one Italian^7^ and one American.^27^ We followed these extensive statistical analyses by functional validation in human iris tissues of the prioritized top gene-level associations. Finally, to gain clinical insights into our findings, we explored the health consequences to individuals carrying high XFS genetic risk in a large biobank with links to electronic health records.

## Materials and Methods

We used an extension of PrediXcan^28^ that uses GWAS summary statistics, S- PrediXcan,^11^ to analyze GWAS summary statistical data from a multi-ethnic GWAS study on XFS.^13^ This dataset consisted of 13,620 XFS cases and 109,837 controls. We also performed PrediXcan on individual-level genetic data from two independent datasets comprising 4127 cases and 9075 controls. The first dataset comprised case and control samples from three cohorts of European ancestry (two from Germany and one from Italy). The second dataset comprised adult patients of European ancestry at Vanderbilt University Medical Center (VUMC) from the local communities surrounding Nashville, TN. The BioVU cases and controls were genotyped on five different Illumina genotyping arrays; Human660W-Quad, HumanOmni1-Quad, Infinium Omni5-4, OmniExpress-8v1-2-B and Infinium Multi-Ethnic Global-8 (MEGA). The data was processed using established GWAS quality control procedures^8^, and imputed on the Michigan Imputation server. Details on how subject selection for BioVU data and genotyping was performed is found in extended materials and methods section (**Supplemental Information**).

### Statistical Analysis

We used the gene-based method, PrediXcan, that provides a framework for correlating imputed gene expression with phenotype.^9^ Gene expression prediction models for 48 different human tissues were trained using GTEx v7 data, subsampled to use only the European ancestry samples. Models with non-zero weights that met a set significance criterion (r > 0.10, q < 0.05) were retained.^27^ Given the lack of eye tissue in the GTEx data, we performed PrediXcan analysis in all available tissues to leverage the shared regulatory architecture of gene expression across tissues.^29^ We referred to the association analysis in each tissue between predicted expression and XFS as “single-tissue analysis.” Because XFS is considered a systemic disorder, we also aggregated evidence across the different tissues to improve our ability to prioritize genes relative to a single unrelated tissue. We determined the joint effects of gene expression variation predicted across all 48 tissues using the Multi-Tissue PrediXcan (MultiXcan), a multivariate regression method that integrates evidence across multiple tissues taking into account the correlation between the tissues.^30,31^ We refer to this association analysis as “multi-tissue analysis.”

We used S-PrediXcan^28^ to analyze all GWAS summary statistic data from the multiethnic study of Aung, *et al.*^13^. Since the summary-based method has been shown to be conservative and tends to underestimate significance in cases where there is some linkage disequilibrium-structure mismatch between reference and study cohorts,^28^ we retained and reported S-PrediXcan results that had a univariate S-PrediXcan *P*<0.0001. We used Bonferroni adjustment for multiple hypothesis testing. Genome-wide significance for a gene-level association in single-tissue and multi-tissue PrediXcan analysis were defined as p<2.02e-7 and p<3.02e-6, respectively.

### Conditional Analysis and Linkage Disequilibrium Evaluation

To determine whether multiple association signals within the same locus are due to independent causal genes or statistical artefacts of correlation in measured expression and predicted gene expression for adjacent genes,^32^ we examined the correlation in the gene expression among genome-wide significant genes in the reference GTEx data. We assumed that there is concordance in correlation in measured and predicted gene expressions, but depending on the quality of our predictions, correlation in predicted expression for a pair of genes may be missed. We verified the extent of LD in the 1000 genomes database^33,34^ between variants in the prediction models for significantly associated genes in each tissue.

To measure potential regulatory effects of the two classical *LOXL1* missense variants in our PrediXcan analysis, we excluded them and all the variants in our gene models that were in LD with them (defined as pairwise r^2^ > 0.1) to generate “reduced models”. We predicted gene expression and performed association analysis using reduced models in both the global multiethnic and the European subset for the genes in chromosome 15 region. To assess whether the association signals in the chr15q22-25 region for each tissue are independent of the ‘classical’ *LOXL1* signal, we excluded variants in the prediction models of genes in the region that were in LD (pairwise r^2^ > 0.1) with any variant in the *LOXL1* model. In tissues without a *LOXL1* model (i.e., r^2^ > 0.10, q < 0.05), we excluded variants for chr15q22-25 region genes that were in LD (r2>0.1) with variants in the *STOML1* models. In addition, we excluded variants that were shared between prediction models for genes in the region. In each case, we performed association analysis using the reduced models and compared the results with the original models.

To determine whether additional genes within the region were significantly associated with XFS, independently of the most highly associated genes *(LOXL1* and *STOML1)* identified in the primary analysis, we performed conditional analysis using the actual individual-level genotype data that included our BioVU cohort and a subset of Aung*, et al.* consisting of three European ancestry cohorts. For each tissue with a significant association, the conditional analysis was performed on the gene that was the most statistically significant as identified from the initial PrediXcan analysis. We generated genetically determined expression for each individual in the dataset and then performed association analysis using Genetic Association Analysis Under Complex Survey Sampling (SUGEN: version 8.8)^35^ on the individual imputed gene expression data, including age, sex, first 5 principal components and relatedness in the regression model. A new logistic regression model was then fit to the case-control data by sequentially adjusting for the expression data of the top significant signals as a covariate. We then performed a metaanalysis for the PrediXcan summary statistics from the four datasets. We repeated this procedure until no genes in the region attained our threshold for statistical significance in the tissues tested (<0.05/total # of e-genes x # of tissues tested for each top round of tests).

### Enrichment and pathway analysis

Genes that were predicted to be associated with XFS at genome-wide significance in both single-tissue and multi-tissue analysis, and at nominal significance (p<0.05) in single-tissue analysis were checked for enrichment of particular categories in several databases using the web-based enrichment tools, Enrichr.^36,37^ This was done by using the strongest signal at nominal significance across the 48 tissues for each of the genes analyzed in PrediXcan. Enrichr implements Fisher’s exact test and uses over 100 gene set libraries to compute enrichment.^36^ We also performed rank-based Gene Set Enrichment Analysis (GSEA) using another web-based enrichment tool, 2019 Webgestalt^38–41^ with a more recent database (Gene Ontology January 2019, KEGG Release 88.2, Reactome ver.66 September 2018 and PANTHER v3.6.1 Jan 2018) and the current Reactome database ver. 69 (June 12 2019).^42^ In this case the strongest signal in the PrediXcan result across the 48 tissues for each of the genes analyzed was used. Based on previous studies indicating limitation in accurately quantifying expression effects of variants in highly polymorphic regions,^43,44^ we also performed enrichment analysis after excluding a total of 310 genes in ~6 Mb chromosomes 6 HLA region (hg19 28Mb-34Mb) that encompassed *GPX6 – CUTA* genes (238 genes) and ~2.5Mb chromosome 17 region that encompassed *CCDC43 – NPPEPS* that include the 900 kb inversion common in population of European ancestries (72 genes).

### Quantitative Expression Validation Analysis

#### Human tissues

Human donor eyes used for corneal transplantation with appropriate research consent were obtained from donors of European ancestry. Eyes were processed within 20 hours after death. Informed consent to tissue donation was obtained from the donors or their relatives. The protocol of the study was approved by the Ethics Committee of the Medical Faculty of the Friedrich-Alexander-Universität Erlangen-Nürnberg (No. 4218-CH) and adhered to the tenets of the Declaration of Helsinki for experiments involving human tissues and samples.

For RNA and DNA extractions, 12 donor eyes with manifest XFS syndrome (mean age, 77±9 years) and 19 normal-appearing control eyes without any known ocular disease (mean age, 74±6 years) were used. All individuals who donated the XFS tissues were previously confirmed XFS patients through routine ophthalmologic examination after pupillary dilation. The presence of characteristic XFS material deposits in manifest disease was assessed by macroscopic inspection of anterior segment structures and confirmed by electron microscopic analysis of small tissue sectors. Iris tissues were prepared under a dissecting microscope and rapidly frozen in liquid nitrogen.

#### Real-time PCR

For quantitative real-time PCR, iris tissues were extracted using the Precellys 24 homogenizer and lysing kit (Bertin, Montigny-le-Bretonneux, France) together with the AllPrep DNA/RNA kit (Qiagen, Hilden, Germany) according to the manufacturer’s instructions including an on-column DNaseI digestion step using the RNase-free DNase Set (Qiagen). First-strand cDNA synthesis and PCR reaction was performed as previously described.^45^ Exon-spanning primers (Eurofins Genomics, Ebersberg, Germany), designed with Primer 3 software (http://bioinfo.ut.ee/primer3/), are summarized in **Suppl. Table S1**. Quantitative real-time PCR was performed using the CFX Connect thermal cycler and software (Bio-Rad Laboratories, München, Germany). Probes were run in parallel and analysed with the ΔΔCt method. Averaged data represent at least three biological replicates. Unique binding was determined with UCSC BLAST search (https://genome.ucsc.edu/) and amplification specificity was checked using melt curve, agarose gel and sequence analyses with the Prism 3100 DNA-sequencer (Applied Biosystems, Foster City, CA). For normalization of gene expression levels, mRNA ratios relative to the house-keeping gene GAPDH were calculated.

Group comparisons were performed using a Mann-Whitney U test using SPSS v.20 software (IBM, Ehningen, Germany). *P* < 0.05 was considered statistically significant.

### Testing for Comorbidity/Pleiotropy

To determine the comprehensive health consequences of high genetic risk to XFS, we performed a phenome-wide association study (PheWAS).^46^ First, we examined the comorbidity of other phecodes with XFS (365.5 – ICD9 365.52/ICD10 H40.14xx) in a total of 752,024 individuals in the VUMC EHR (418,371 females and 333,653 males), by performing logistic regression analysis conditioned on gender, age and the self-reported ancestry as covariates in the regression model. For this analysis we used a total of 600,107 European, 103,209 African, 12,411 Asian and 36,297 other ancestry patients, of which 222 were uncurated XFS cases (coded as 1) and the rest controls. To determine other health consequences of high genetic risk to XFS, we performed a PheWAS analysis^30^ (n=52,251) on the polygenic risk score generated from the Aung *et al’s*^13^ XFS global dataset against patients genotyped on Illumina Mega-array chip in BioVU with about 18k ICD-9 /ICD-10 codes, accounting for age, gender, and the first 5 principal components.

## Results

### PrediXcan Analysis

We performed single-tissue PrediXcan analysis of the global multi-ethnic GWAS (13,620 XFS cases and 109,837 controls) summary data, identifying 23 genes (defined as signals with *P*<2.02 x 10^-7^ after Bonferroni corrections) on chromosome 15: *CYP1A2, CYP1A1, STOML1, LOXL1, ISLR2, RPP25, INSYN, ISLR, STRA6, CD276, NEO1, ARID3B, COX5A, PML, CPLX3, LMAN1L, UBL7, MPI, CLK3, CSK, SEMA7A, TBC1D21,* and *NPTN*(**Figure 1a, Suppl. Table S2**).

**Figure 1b:**
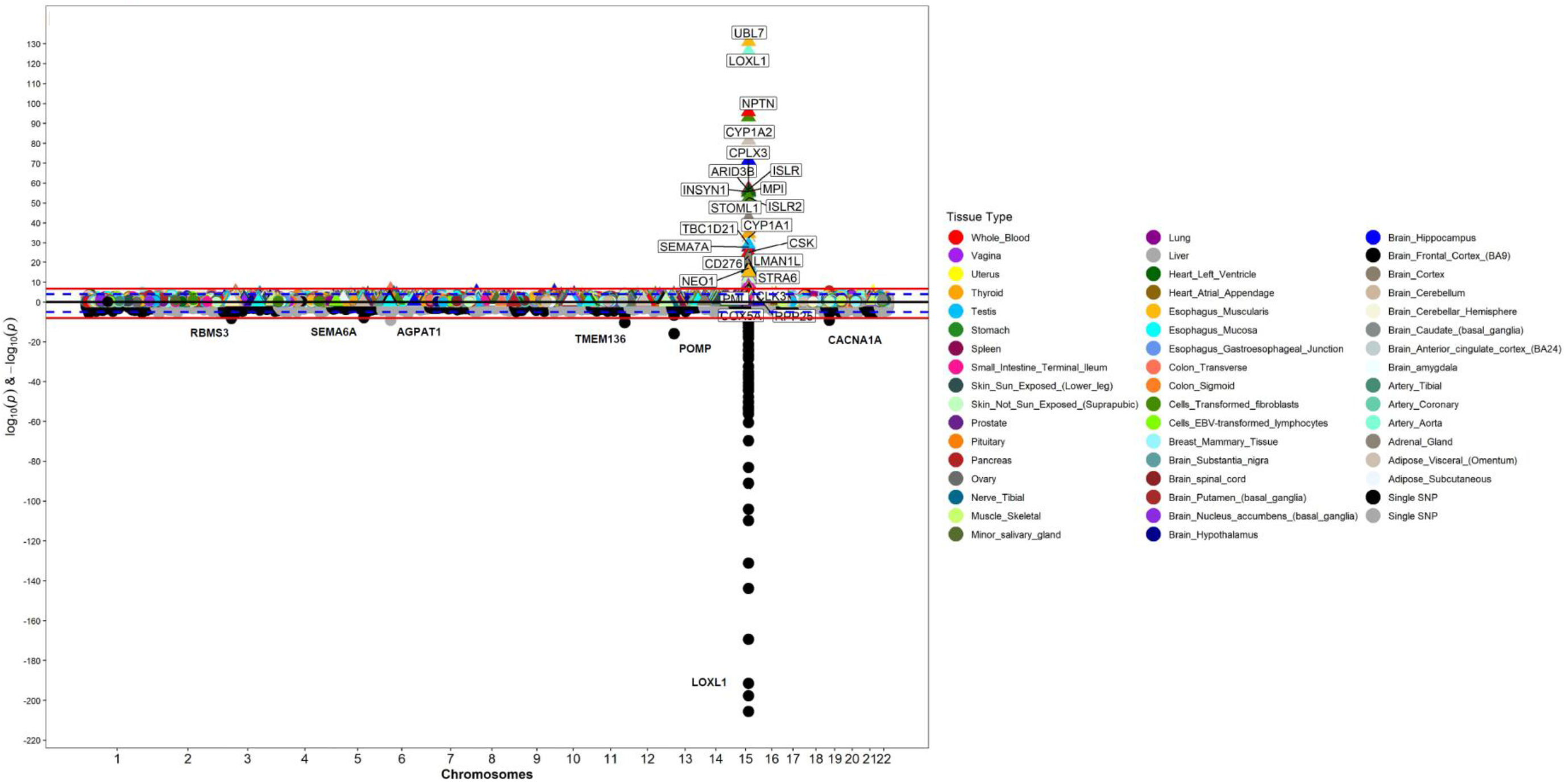
Manhattan plot for GWAS meta-analysis and PrediXcan analysis of the genotyping data for XFS. The lower half of the plot is for the XFS meta-analysis summary statistics data Aung *et al.,* 2017, while the upper half of the plot shows results from PrediXcan analysis for 48 GTEx tissues. On the X axis is plot of variant/gene associations along the chromosomes, while Y axis represent the significance levels for the associations. The legend for PrediXcan analysis on the 48 GTEx tissues, a color for each tissue, is on the right. For both plots the blue dotted line is the “suggestive” genome-wide significant threshold (p<1e-4), while the red line is the genome-wide significant threshold. On the lower plot, the gene labels are for genes reported/mapped to genome-wide significant signals in GWAS result, while in the upper plot is for genes that are associated at genome-wide significant threshold. For genes associated with XFS at genome-wide threshold in more than one tissues, only the tissue with lowest p-value is labeled. The GWAS plot has been truncated to p<1e-220 for clarity.

To determine the joint effects of gene expression variation predicted across all 48 tissues analyzed, we performed a multivariate regression multi-tissue analysis. Each of the 23 associations from the single-tissue PrediXcan analysis remained significant in multi-tissue analysis (**Suppl. Table S3**). Additionally, five genes within the same region that were associated with XFS at subgenome-wide significance in the single-tissue analysis (p<3.02e-6) were associated in the multi-tissue analysis: *ADPGK* (p=7.32E-07), *CYP11A1* (p=1.36E-16), *HEXA* (p=1.03E-06), *PARP6* (p=1.82E-06), *SCAMP2* (p=1.65E-10). All genes mapped to chromosome region 15q22-25, spanning ~3 Megabases (**Figure 1b)**.

**Figure 1b:**
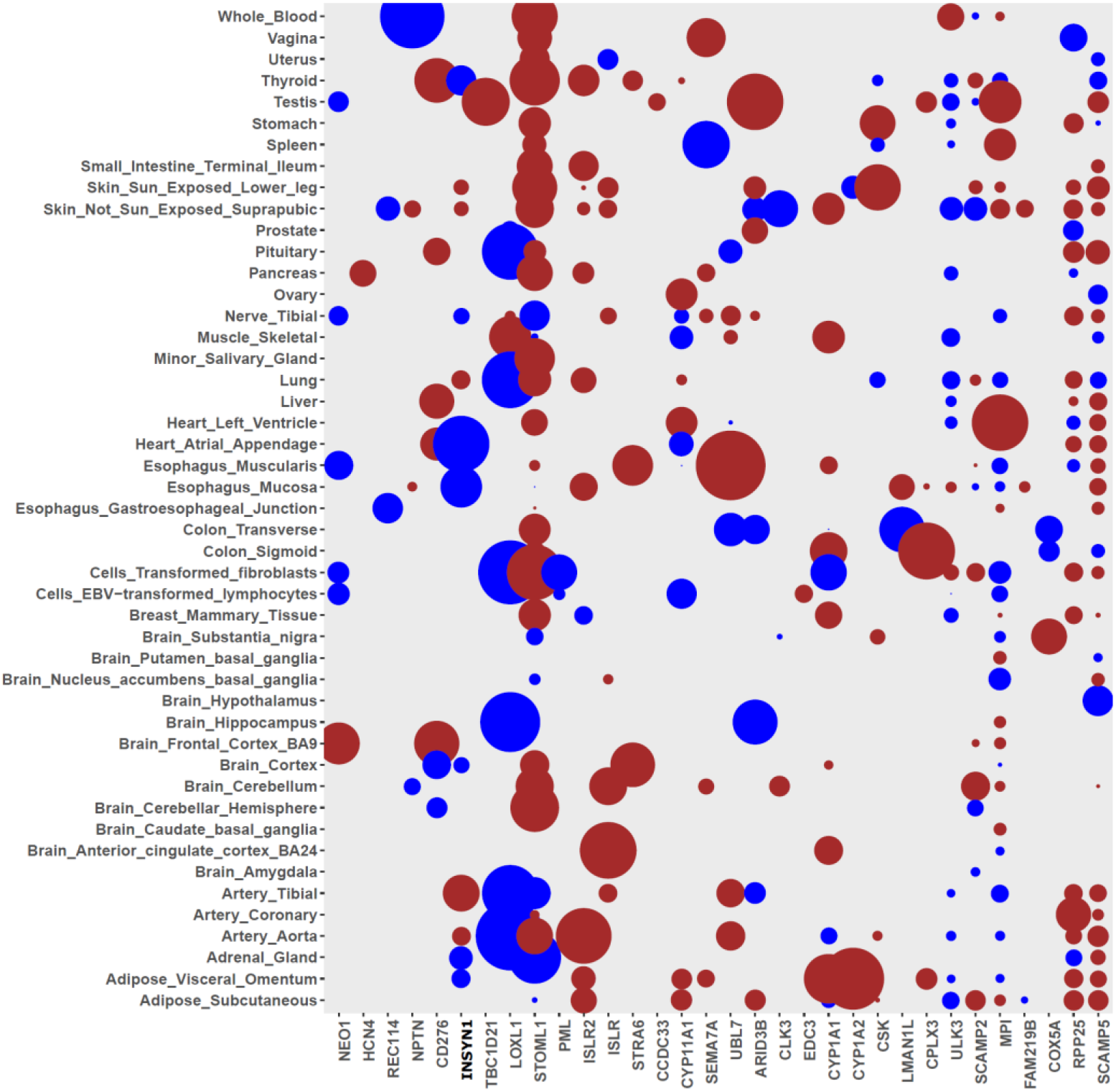
Chromosome 15q22-25 region genes that show significant association with XFS. The size of the balloon for each gene-tissue association is proportional to -log10_pvalue_ and color corresponds to the predicted direction of expression changes: dark-red and blue for increased and decreased expression changes, respectively. Only four genes (EDC3, ULK3, HCN4 & FAM219B) in the whole region were not associated with XFS.

Seven additional genes located on chromosomes 1 *(LGR6* p=2.20E-06; *SDHB* p=8.07E- 08), 6 *(PRRT1* p=9.10E-07), 8 *(PRSS55* p=4.18E-13), 10 *(CDH23* p=1.86E-07; *PITRM1* p=8.45E-12) and 19 *(CALM3* p=2.60E-07), were significantly associated in the multi-tissues analysis (**Figure 1a, Suppl. Table S3**). All seven signals mapped to genomic regions harboring GWAS SNP variants showing subgenome-wide significance with XFS risk, except for PRRT1, which corresponds to the AGPAT1 locus.^13^ The data indicates that combining information across variants in genes and then across tissue expression improves the power to identify additional XFS-associated loci.

To ensure that the association observed at the 23 genes from the larger multi-ethnic dataset was not an artefact of population structure, we confirmed the signals in a subset of European ancestry individuals (**Materials and Methods**). Twelve out of the 23 genes in chr15q22-25 region (ten in the single-tissue analysis and two in the multi-tissue analysis) remained genome-wide significant in this European ancestry analysis, whilst the remaining 11 genes remain nominally associated (p<0.05) (**Table 1, Suppl. Figure S3, Suppl. Table S4, S5, S6**).

**Table 1:**
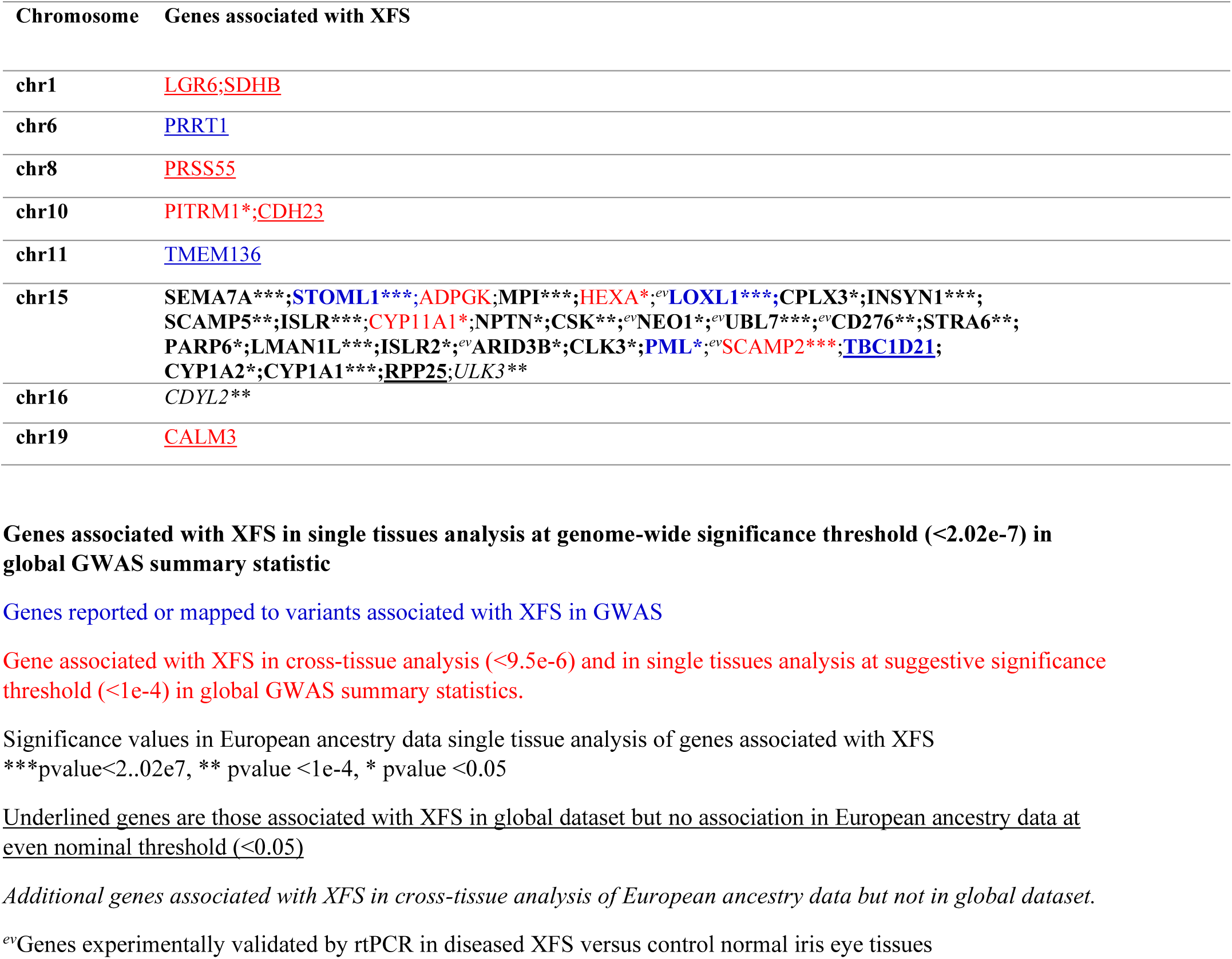
Confirmation of PrediXcan analysis of global dataset in European ancestry individuals

### Correlated Expression Among Significant Genes

To determine whether the 23 observed association gene signals were artefacts of LD contamination, we performed extensive additional analyses. We calculated the pair-wise correlation in measured expression among the significant genes, using the reference GTEx panel. We checked the relationship between expression correlation for each of the chr15p22-25 genes with *LOXL1* and *STOML1* and the PrediXcan associations for the two genes in each tissue. We made two important observations from this analysis. First, there was a significant correlation between the correlation of measured gene expression of the other genes in chr15p22-25 with *LOXL1* or *STOML1* and the gene-level associations with XFS in most tissues **(Suppl. Table S7a, Suppl. Figure S4a)**. Secondly, there is substantial correlation between *STOML1* and *LOXL1* (r^2^ = 0.67, p=0.009) (**Suppl. Table S7a, Suppl. Figure S4a)**. These results indicate that the associations by one of the genes might be due to LD contamination or the presence of shared variants in the prediction models of the two genes (**Suppl. Table S8**).

To dissect the potential source of LD contamination in the PrediXcan analysis, we looked into the effect of the two GWAS missense variants implicated in XFS that have mostly been linked to *LOXL1* and shown to play regulatory roles,^23^ followed by the effect of *LOXL1* and *STOML1* signals on chr15q22-25 region observed associations in each tissue. We also determined the effect of shared variants between prediction models for the genes in the region.

We modified our prediction models by excluding: i) rs3825942 missense variant, ii) rs4886776 intronic variant, which is at near perfect LD (pair-wise, *r^2^*=0.982) with the rs1048661 missense variant, and iii) all the variants in our gene models that were in LD with the two variants *(r^2^* > 0.1). The two missense variants had wide ranging effect on the genetically predicted expression of many chr15q22-25 region genes, with the largest effect on *LOXL1.* The strength of the association signals diminished in six of the nine tissues for which we had the gene’s predicted expression. Association signals in three of these tissues fell below genome-wide threshold in the global dataset **(Suppl. Table S7b)**. In addition, association signals for seven additional genes in the region besides *STOML1* lost genome-wide significance; *CD276* (2 tissues)*, COX5A, CYP1A1, LMAN1L, MPI, SCAMP2 and TBC1D21.*

Interestingly, association signals for eight genes were strengthened, four of which attained genome-wide significance threshold in reduced models: *INSYN1, CYP1A1, NPTN* and *LOXL1* **(Suppl. Table S7b)**. These shifts in association strength, i.e., an increase in effect size, seem to be due to the exclusion of select variants **(Suppl. Tables S7c, S7d)**. Moreover, the shifts in association strength are correlated with the excluded variants’ level of LD with the missense variant rs3825942 *(r^2^* = 0.64) **(Suppl. Tables S7c, S7d)**. Notably, the three GWAS variants identified to have effect reversal in South Africans relative to other populations were in high LD with rs3825942 **(Suppl. Table S7g)**.^47^ Our results indicate that the missense variants have enhancing or diminishing effects on the PrediXcan association signals, in chr15q22-25, with XFS, consistent with allele reversal reported for the GWAS variants.^13^

To check whether the association signals in the chr15q22-25 region for each tissue were independent of the ‘sentinel’ *LOXL1* signal, we excluded, from the prediction models, variants that were in LD (*r^2^*>0.1) with any variants in *LOXL1 or STOML1* tissue models. We also excluded variants that were shared between two or more genes in their original prediction models. Seven of the genes that were associated with XFS at genome-wide threshold in their original models showed diminished signals, including four below significance levels: *UBL7, ISLR, LMAN1L* and *COX5A* in reduced models **(Suppl. Table S7e)**. Association signals for *CYP1A1* and *CYP1A2* were slightly diminished in reduced models relative to the original models, but remained at significant genome-wide thresholds **(Suppl. Table S7e)**. However, association signals for six genes strengthened, four of which attained genome-wide association significance levels in the reduced models: *INSYN1, CLK3, CYP1A1* and *NEO1* **(Suppl. Table S7e).** These shifts in association strength seem to be due to few variants that are either in LD with variants in *LOXL1* and *STOML1* models or are shared with other genes’ models **(Suppl. Tables S7e, S7f)**. However, these variants causing the shifts in association signals upon exclusion from the models, were not in LD with the missense rs3825942 variant **(Suppl. Table S7g).** This indicates that there are signals of allele reversal independent of the known missense variants in *LOXL1*.

Excluding variants that are in LD with SNPs in the *LOXL1/STOML1* models did not have any effect on the association signals for six genes that were associated with XFS at genome-wide threshold in the original models: *CSK, STRA6, CD276, ARID3B, MPI & TBC1D21,* with the latter three in testis, for which we had no models for both *LOXL1* and *STOML1* **(Suppl. Table S7e)**. The results indicate that some of the observed signals were artefacts of LD contamination from *LOXL1* and *STOML1 (ISLR, LMAN1L* and *COX5A),* while some of the signals were masked in the original models *(INSYN1, CLK3, CYP1A1* and *NEO1).* There was inconsistent result for *UBL7,* where there was no effect in its association signal in a tissue, enhanced effect in another tissue, and diminished signal in two other tissues, one of which went below the genomewide threshold, albeit the reduced model had only a single variant in the prediction **(Suppl. Table S7e).**

### Conditional Analysis

Conditional analysis was performed in tissues with any genome-wide significant chr15p22-25 region gene signals against the predicted gene expression for the strongest observed signals in the European subset. As in the global dataset, the strongest signals in the European dataset were at *LOXL1* **(Table 1, Suppl. Figure S3)**. In all nine of the 48 tissues with *LOXL1* predicted expression, only the *STOML1* gene showed a significant association signal (in addition to *LOXL1)* **(Table 1, Suppl. Figure S3)**. After conditioning on *LOXL1* in these tissues, the *STOML1* signal disappeared, but association signals at *SCAMP2* and *INSYN1* were observed in artery-aorta and lung tissues, respectively **(Suppl. Table S8, Figure 2a, 2b)**. This indicated that the association of *STOML1* with XFS is an artefact of a strong *LOXL1* signal, consistent with *LOXL1* being the true signal and *STOML1* a proxy signal. In addition, association signals for *SCAMP2* and *INSYN1* were masked by the *LOXL1* signal.

**Figure 2a, 2b:**
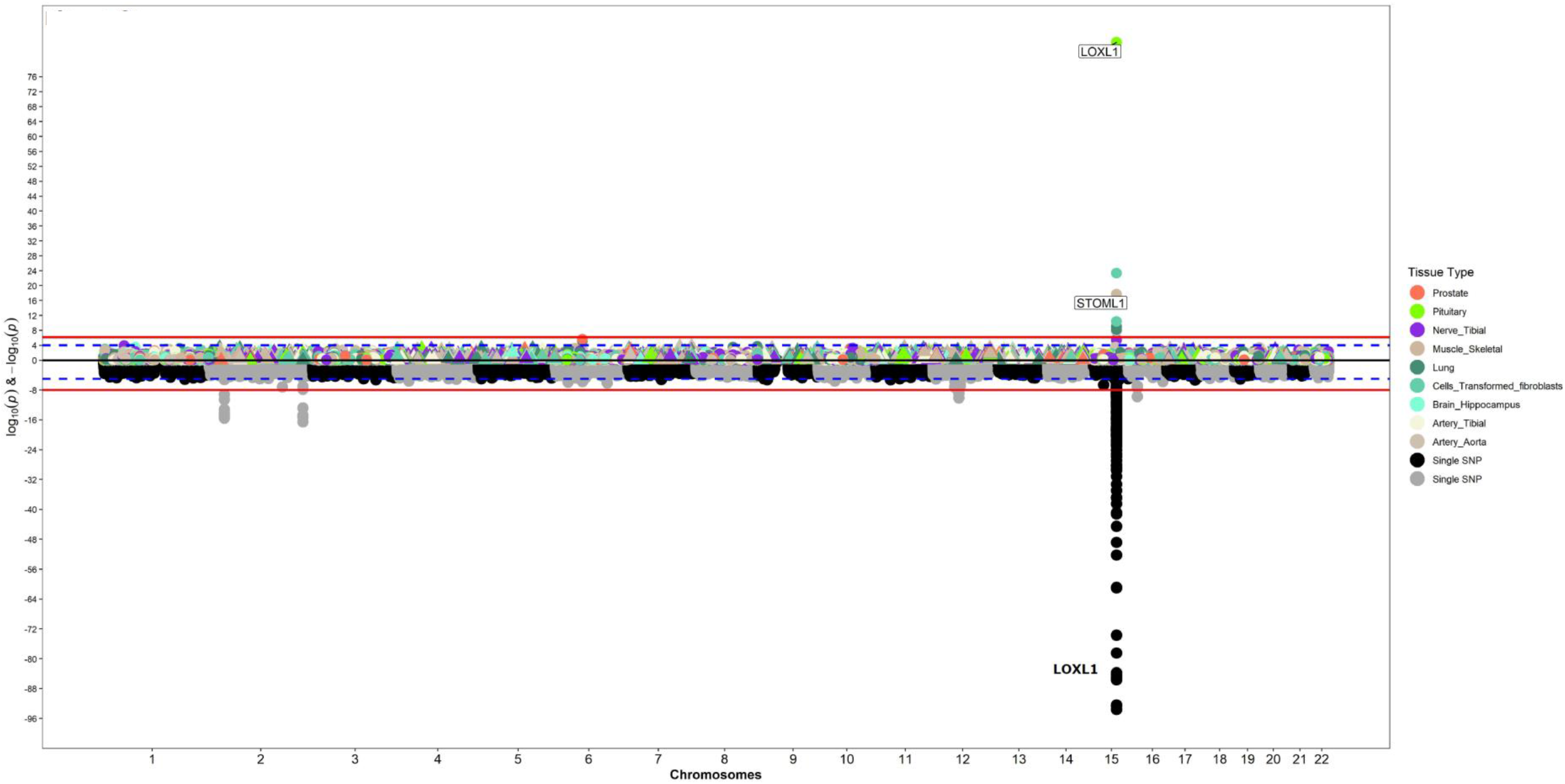
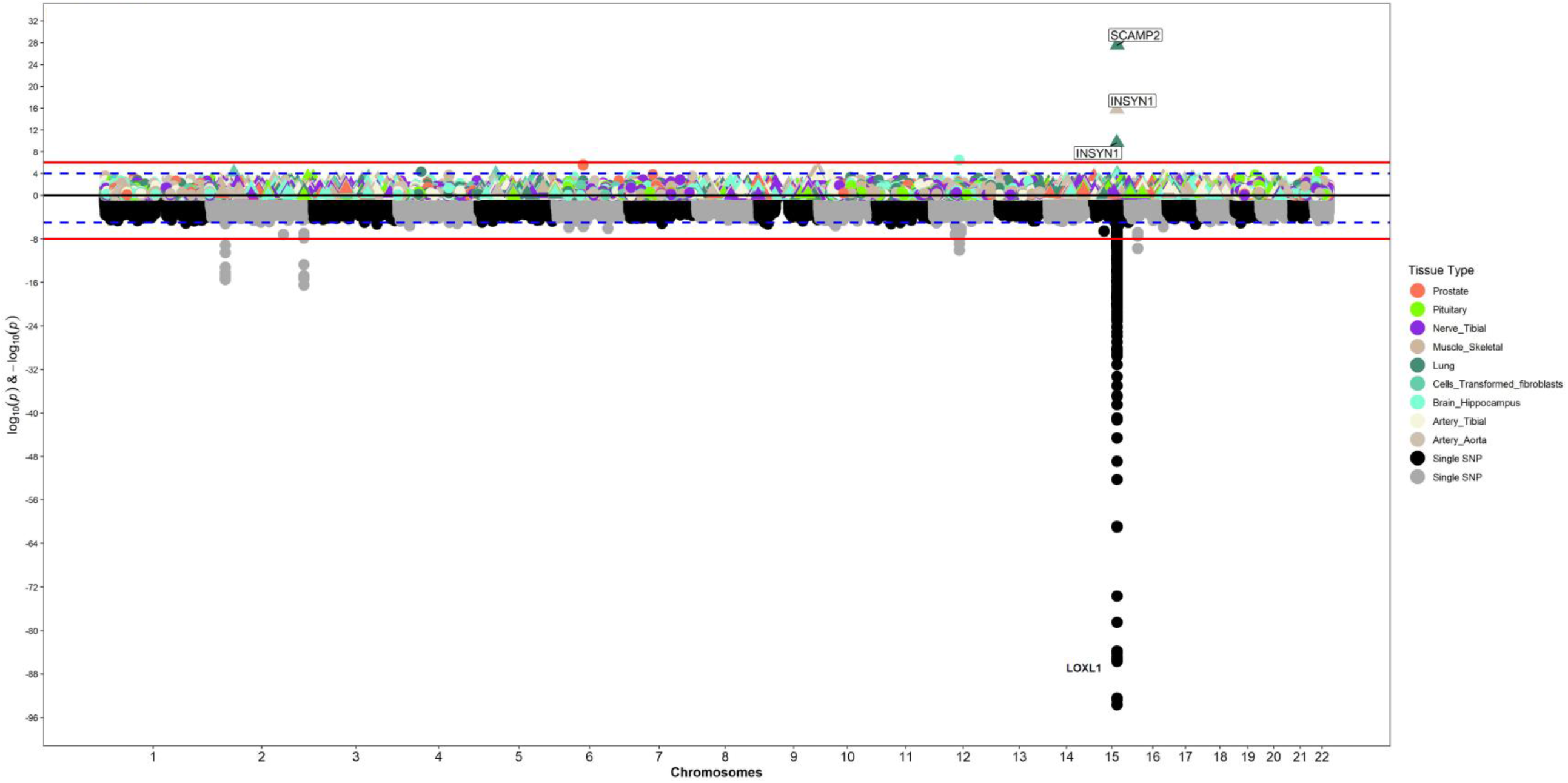
Conditional analysis to prioritize XFS associated genes: Manhattan plot for PrediXcan analysis of European ancestry individuals in tissues with predicted gene expression for **a)** LOXL1 and **b)** conditioned on LOXL1 predicted gene expressions In each case on the X axis is plot of variant/gene associations along the chromosomes, while Y axis represent the significance levels for the associations. The legend for PrediXcan analysis on the GTEx tissues, a color for each tissue, is on the right. For both plots the blue dotted line is the “suggestive” genome-wide significant threshold (p<1e-4), while the red line is the genome-wide significant threshold. On the lower plot, the gene labels are for genes reported/mapped to genome-wide significant signals in GWAS result, while in the upper plot is for genes that are associated at genome-wide significant threshold. For genes associated with XFS at genome-wide threshold in more than one tissues, only the tissue with lowest p-value is labeled.

In 17 tissues with *STOML1* predicted gene expression, we observed significant association signals for 8 other genes (in addition to *STOML1*) **(Suppl. Table S8, Figure 2c, 2d)**. After conditioning on *STOML1* predicted gene expression, associations with four genes *(CYP1A1, INSYN, LOXL1, SCAMP2)* remained, while association with four other genes *(ISLR, LMAN1L, MPI & SEMA7A)* disappeared, consistent with the role of gene expression correlation in our ability to ascertain true association **(Figures 2e-2g)**. In addition, associations with five more genes *(ARID3B, CPLX3, CYP1A2, PML & UBL7)* attained genome-wide significance after the conditional analysis.

**Figure 2c-2g:**
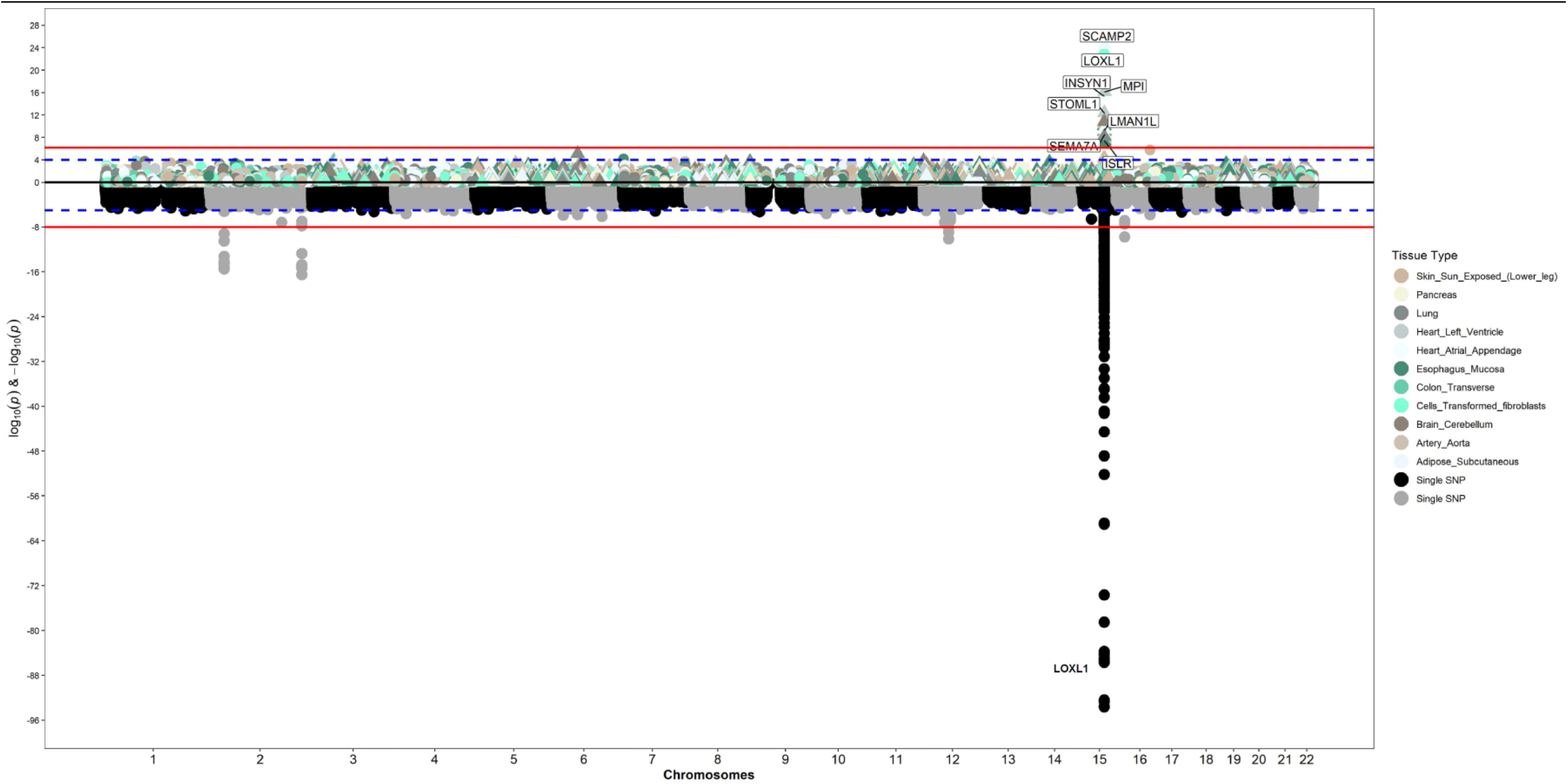
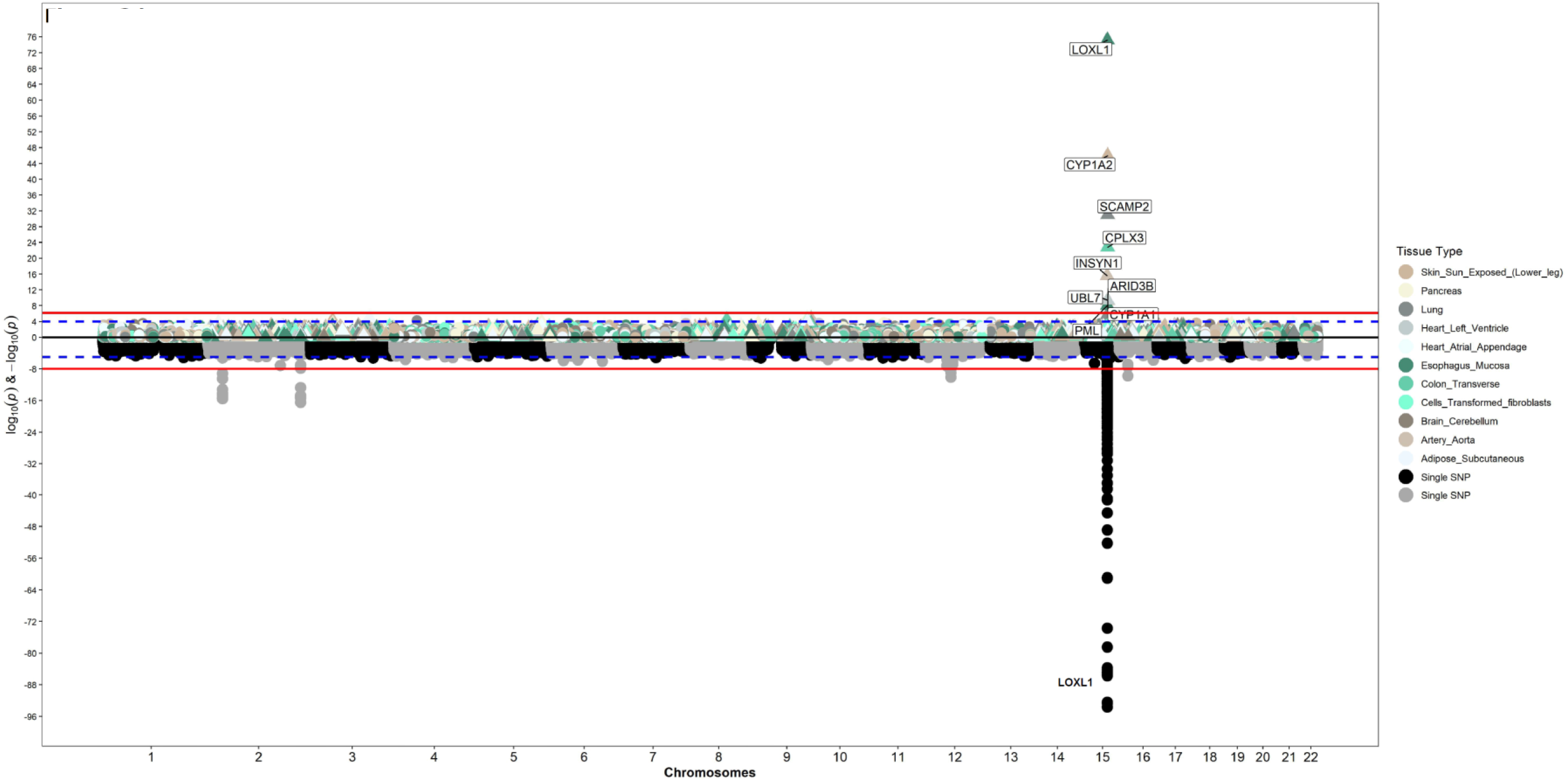
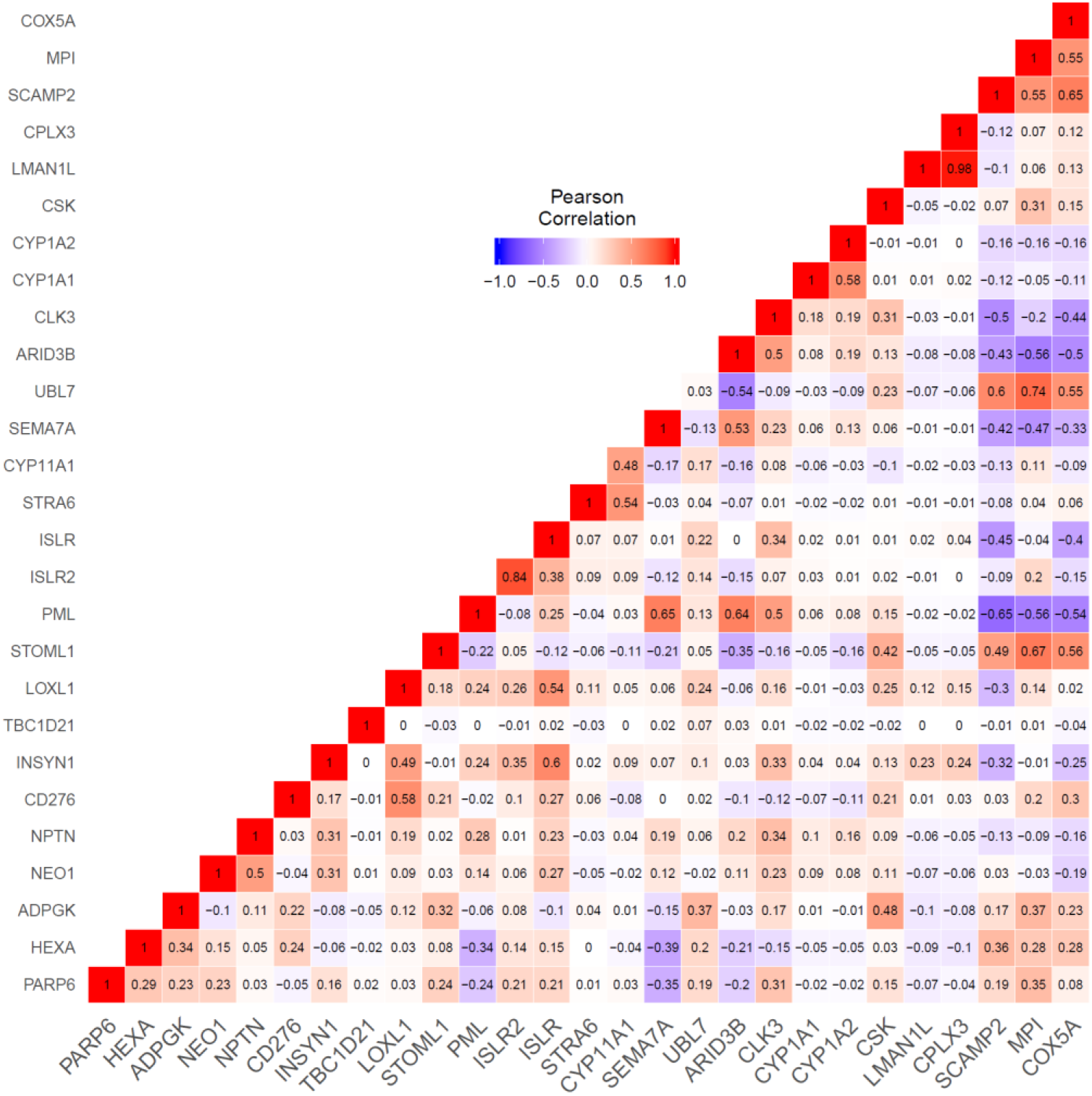
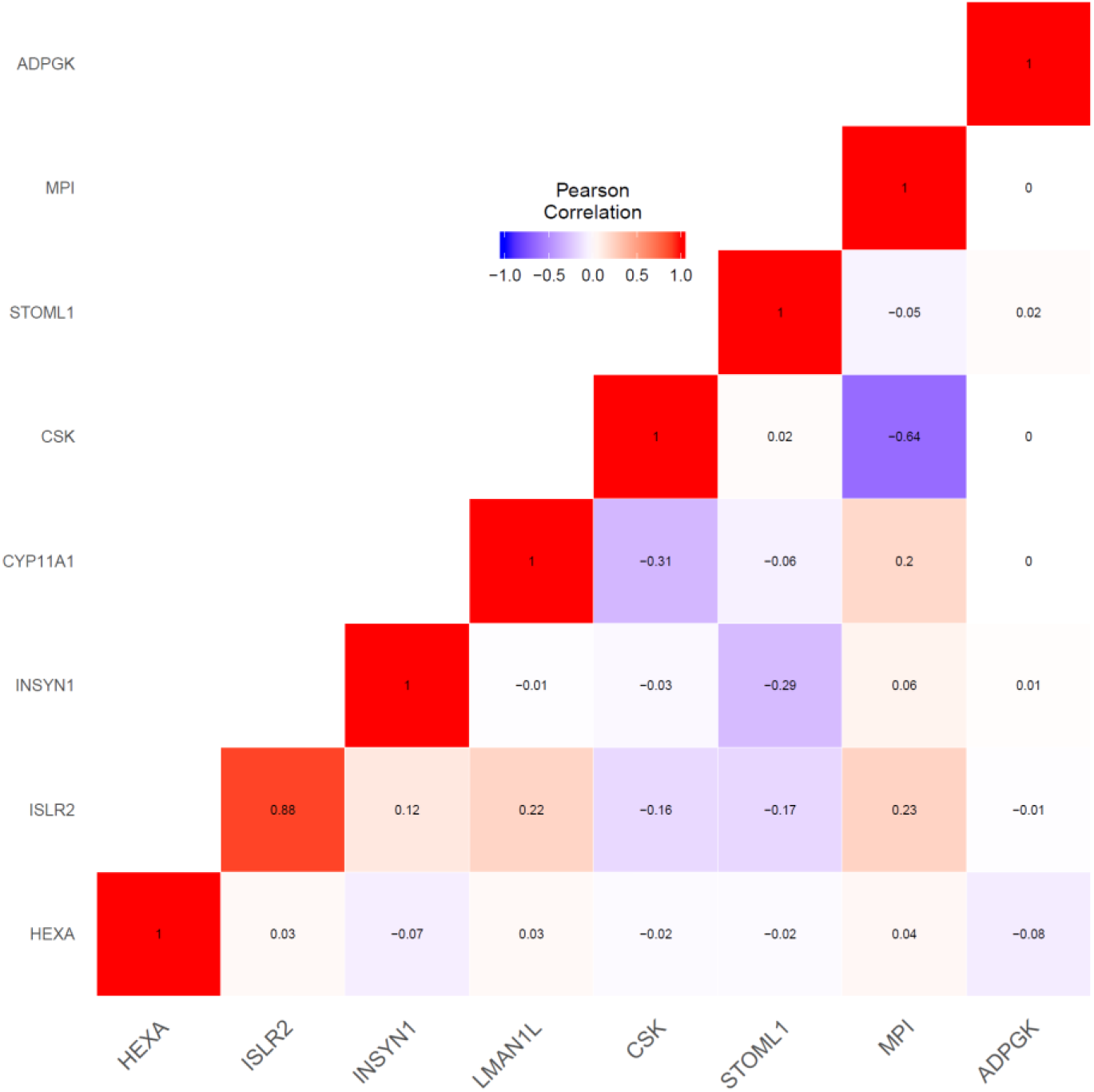
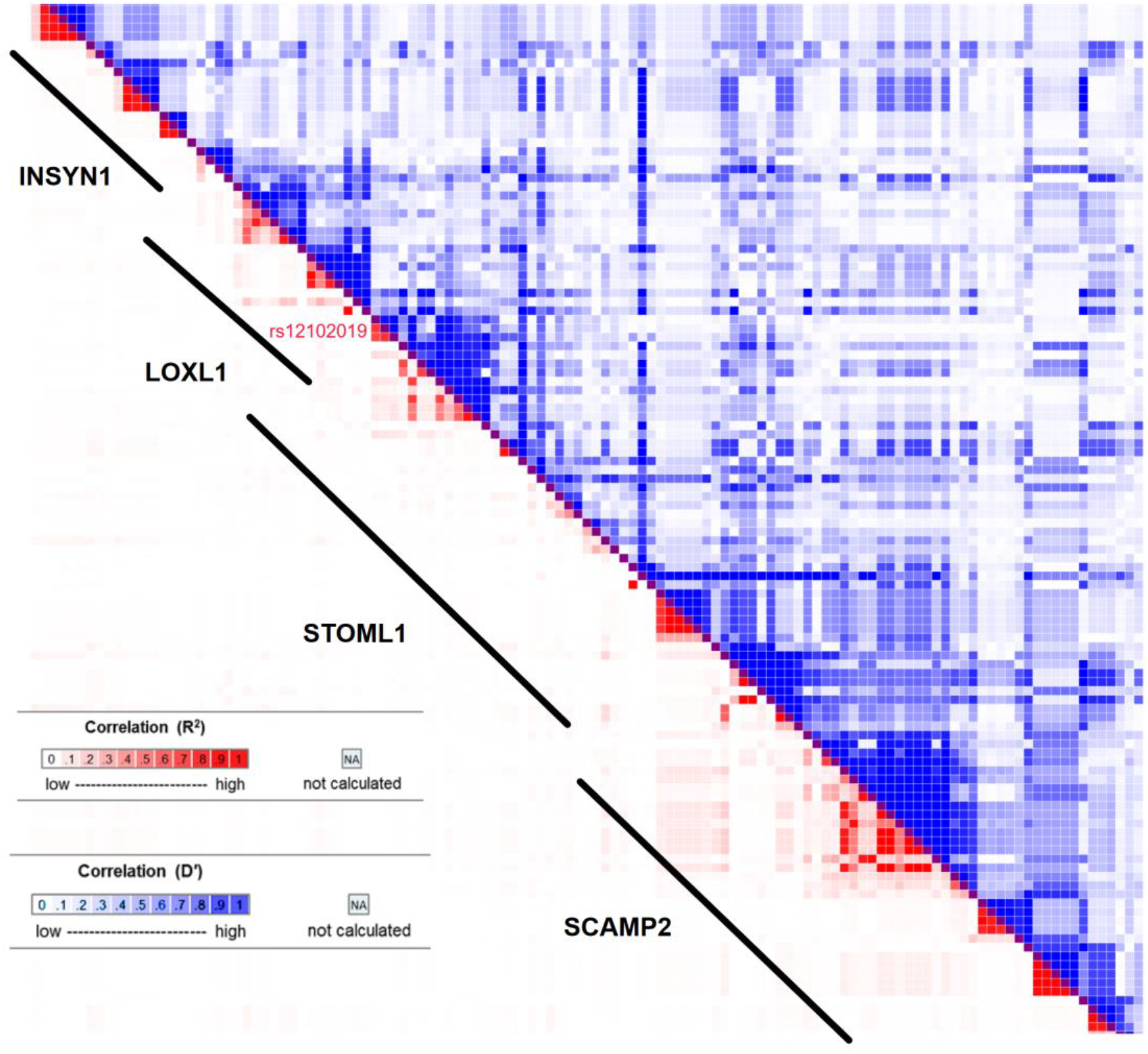
Conditional analysis to prioritize XFS associated genes: Manhattan plot for PrediXcan analysis of European ancestry individuals in tissues with predicted gene expression for STOML1 (**2c)** and plot of analysis conditioned on STOML1 predicted gene expressions (**2d). Figure 2e)** correlation in gene expression in for genes in chr15q22-25 in lung tissue for i) reference GTEx data **f)** predicted gene expression in BioVU cohort. In each case on the X axis is plot of variant/gene associations along the chromosomes, while Y axis represent the significance levels for the associations. The legend for PrediXcan analysis on the GTEx tissues, a color for each tissue, is on the right. For both plots the blue dotted line is the “suggestive” genome-wide significant threshold (p<1e-4), while the red line is the genome-wide significant threshold. On the lower plot, the gene labels are for genes reported/mapped to genome-wide significant signals in GWAS result, while in the upper plot is for genes that are associated at genome-wide significant threshold. For genes associated with XFS at genome-wide threshold in more than one tissues, only the tissue with lowest p-value is labeled. **g)** linkage disequilibrium between variants in prediction models for LOXL1 and other chr15q22-25 genes associated with XFS in lung tissue based on pairwise r^2^ and D’ parameters. Relative genome location for variants in each gene models are roughly demarcated by diagonal lines next to gene symbols. Proximate location for the variant shared between LOXL1 and STOML1, rs12102019 is labelled.

Overall, conditional PrediXcan analysis of genetic signals in the chromosome 15 region in the European dataset in a limited number of tissues was mostly consistent with PrediXcan analysis using the reduced models above. The analysis confirms the associations for *LOXL1, ARID3B, CPLX3, CYP1A1, CYP1A2, INSYN1, NEO1, PML, SCAMP2,* and *UBL7,* all of which, except for *INSYN1,* have been shown to be highly expressed in eye tissues^48^ (**Suppl. Figure S5**). However, *INSYN1* has enhanced expression in brain tissues.^49,50^ Collectively, these results suggest that some of the identified gene-level association signals between XFS and genetically imputed expression were driven by correlation to the strong *LOXL1* and its “proxy” *STOML1* signal.

### Enrichment and pathway analysis

Genes at genome-wide significance (p<2.02e-7) and nominal significance (p<0.05) were evaluated for enrichment of known pathways, using Enrichr.^36,37^ Genes at genome-wide significance were enriched for genes reported for, or mapped to, GWAS variants implicated in several caffeine-related (coffee and caffeine consumption, and caffeine metabolism^51–53^) and blood pressure^54^ traits. The enrichment for coffee consumption is replicated for the larger gene set that is associated with XFS at nominal significance.^55^ Some of these genes, *CYP1A1* and *CYP1A2*,^56^ are involved in fatty acid oxidation and estrogen receptor pathways. In addition, these two genes are also observed in the Reactome enrichment of protectin synthesis **(Table 2, Suppl. Table S9)**.

**Table 2:** Enrichment analysis of genes associated with XFS

| Tool | Database | enrichment | Name | #found<sup>1</sup> | #total<sup>2</sup> | Adj-p-values/FDR<sup>a</sup> |
| --- | --- | --- | --- | --- | --- | --- |
| **Reactome** | reactome | pathway | Endosomal/Vacuolar pathway | 59 | 82 | 1.83E-07 |
| **GSEA** | reactome | pathway | Crosslinking of collagen fibrils | 5 | 8 | 0.031 |
|  | Wikipathway | pathways | Aryl Hydrocarbon Receptor Pathway | 6 | 12 | 0.013 |
|  |  |  | Aryl Hydrocarbon Receptor | 6 | 13 | 0.017 |
|  |  |  | Oxidation by Cytochrome P450 | 3 | 15 | 0.018 |
|  |  |  | Fatty Acid Omega Oxidation | 5 | 7 | 0.047 |
|  | DrugBank | Drugs | Clomiphene | 3 | 5 | 0.006 |
|  |  |  | Ketoconazole | 3 | 7 | 0.006 |
|  |  |  | Quinidine | 2 | 9 | 0.007 |
|  |  |  | Acetaminophen | 2 | 7 | 0.007 |
|  |  |  | Estradiol | 2 | 5 | 0.007 |
|  |  |  | Estradiol acetate | 2 | 5 | 0.007 |
|  |  |  | Estradiol benzoate | 2 | 5 | 0.007 |
|  |  |  | Estradiol cypionate | 2 | 5 | 0.007 |
|  |  |  | Estradiol dienanthate | 2 | 5 | 0.007 |
|  |  |  | Estradiol valerate | 2 | 5 | 0.007 |
| **Enrichr** | Jensen Diseases<sup>b</sup> | Disease | Rheumatoid_arthritis | 119 | 310 | 8.32E-08 |
|  |  |  | Type_1_diabetes_mellitus | 68 | 158 | 2.69E-06 |
|  |  |  | Carcinoma | 2619 | 11318 | 0.013 |
|  |  |  | Vitiligo | 28 | 63 | 0.029 |
| **Enrichr** | GWAS Catalog<sup>c</sup> | traits | Caffeine consumption | 11 | 14 | 0.019 |
| **Enrichr** | DSigDB<sup>d</sup> | Drugs | Cyclosporin A_CTD_00007121 | 1258 | 4826 | 9.66E-11 |
|  |  |  | Valproic acid_CTD_00006977 | 2041 | 8313 | 1.75E-09 |
|  |  |  | Copper sulfate_CTD_00007279 | 1508 | 6017 | 3.04E-08 |
|  |  |  | Quercetin_CTD_00006679 | 812 | 3159 | 7.82E-05 |
|  |  |  | acetaminophen_CTD_00005295 | 1017 | 4136 | 0.007 |
|  |  |  | Aflatoxin B1_CTD_00007128 | 773 | 3082 | 0.006 |
|  |  |  | (-)-Epigallocatechin gallate_CTD_00002033 | 546 | 2115 | 0.005 |
|  |  |  | Potassium chromate_CTD_00001284 | 491 | 1898 | 0.011 |
|  |  |  | Methamphetamine_CTD_00006286 | 40 | 102 | 0.030 |
|  |  |  | Methyl Methanesulfonate_CTD_00006307 | 940 | 3865 | 0.047 |

<sup>1</sup>number of genes that are associated with XFS that belong gene set under test
<sup>2</sup>total number of genes in gene set being tested
<sup>a</sup>FDR fro Reactome and GSEA analysis, Adj-p.values for Enrichr
<sup>b</sup>Jensen Diseases – enrichment for genes associated with diseases in gene-diseases association mined from literature (https://diseases.jensenlab.org/).
<sup>c</sup>GWAS catalog – enrichment for genes reported for GWAS variants in variant-traits association (https://www.ebi.ac.uk/gwas/)
<sup>d</sup>DSigDB – enrichment for genes that are targets for a compound/drug (http://tanlab.ucdenver.edu/DSigDB)

Our gene set is also enriched for genes associated with carcinoma and three inflammatory conditions: rheumatoid arthritis, Type 1 diabetes, vitiligo in Jensen Diseases, a database that integrates evidence on disease-gene associations from automatic text mining, manually curated literature, cancer mutation data, and GWAS (https://diseases.jensenlab.org/).

We further analyzed our gene list against compounds in Drug Signatures Database (DSigDB, http://tanlab.ucdenver.edu/DSigDB), a gene set resource that relates drugs/compounds and their target genes. Our gene set is enriched for genes that are targets of cyclosporin A (p=9.66E-11), and genes that are targets for compounds that are either 1) carcinogenic: Aflatoxin B1, potassium chromate, methyl methanesulfonate and copper sulfate, 2) neuroactive: valproic acid and methamphetamine, 3) neuroprotective: quercetin and epigallocatechin gallate, or 4) analgesic: acetaminophen (**Table 2, Suppl. Table S9**). Cyclosporin A is an immunosuppressant taken to treat rheumatoid arthritis and other autoimmune conditions, while quercetin and acetaminophen have been shown to have anti-inflammatory effects.^57,58^

Analysis in Gene Set Enrichment Analysis (GSEA) using a ranked association gene list based on effect sizes confirmed some of the enrichment observations using Enrichr. GSEA besides replicating enrichment for acetaminophen, showed enrichment for: 1) six synthetic estrogens, 2) estrogen regulators (Clomifene), 3) antiarrhythmic (quinidine), and 4) an antifungal (ketaconazole) **(Table 2, Suppl. Table S9).** The gene set was also enriched for genes that were associated with the collagen fibril crosslinking (FDR=0.0313) Reactome pathway. Analysis of the gene list in relation to the latest Reactome library (https://reactome.org/) returned significant enrichment for the endosomal-vacuolar pathway (p=8.14E-11), an enrichment that was replicated in gene sets that were predicted to be downregulated (p=3.24E-8). Our results broadly recapitulated results above, even after excluding genes in HLA and chr17 inversion regions from the enrichment analysis of the gene set (p<0.05) **(Table 2, Suppl. Table S9)**

### Quantitative Expression Validation Analysis

Expression levels of *ARID3B, CD276, INSYN1, LOXL1, NEO1, SCAMP2, STOML1* and *UBL7* were measured in XFS and control eye tissues. All transcript levels were found to be decreased in iris tissues obtained from XFS patients compared to control samples, with significant differences for *ARID3B, CD276, LOXL1, NEO1, SCAMP2* and *UBL7* (*p*<0.05) **(Figure 3)**. *INSYN1* and *STOML1* were not significantly downregulated in diseased eyes relative to normal eyes in validation analysis. *STOML1* is the closest gene to and potentially proxy for *LOXL1* among those that show association in our PrediXcan results within the chr15q22-25 region. We included it as a negative control in the validation analysis, while *LOXL1* was a positive control considering that it had already been shown to exhibit pattern of downregulation in gene expression in diseased relative to normal tissues.^13^ *CD276* was selected for functional validation in eye tissue despite no significant association with XFS in single-tissue analysis in the European ancestry data because it was significantly associated with XFS in multi-tissue analysis in European data. In addition, it was one of the gene association signals which were not affected by excluding variants that were in LD with *LOXL1/STOML1* model SNPs in the multiethnic global dataset. Overall, our validation results replicate the associations found using the genetically determined gene expression.

**Figure 3:**
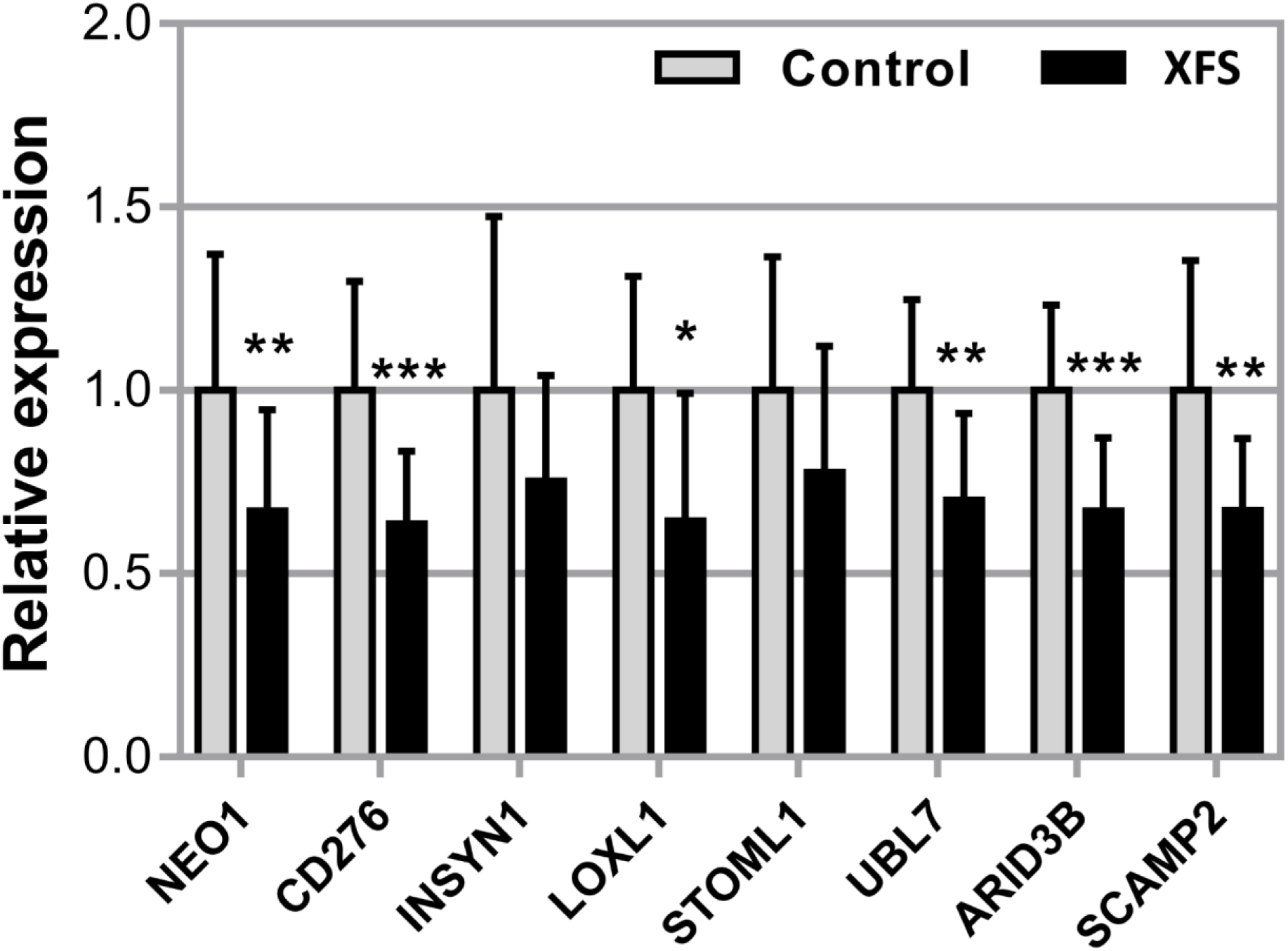
Expression of *NEO1, CD276, INSYN1, LOXL1, STOML1, UBL7, ARID3B* and *SCAMP2* mRNA in iris tissues derived from normal human donors (control) (n=19) and donors with XFS syndrome (n=12) using real-time PCR technology. Expression levels were reduced in XFS specimens compared to control specimens, with significant differences for *NEO1, CD276, LOXL1, UBL7, ARID3B* and *SCAMP2*. The relative expression levels were normalized relative to GAPDH and are represented as mean values ± SD (\**p*<0.05; \*\**p*<0.01, \*\*\**p*<0.001).

### Comorbidity/pleiotropy analysis

To gain further biological insights into the gene associations we observed in our PrediXcan analysis, we performed logistic regression analysis of both XFS ICD9/10 diagnosis and Polygenic Risk Score generated from the multi-ethnic summary data across the BioVU individuals, Vanderbilt University’s electronic health records database linked to genetic information, as the target dataset **(Materials and Methods)**. XFS diagnosis was associated with an increased risk of 96 phenotypes in BioVU, including 12 musculoskeletal phenotypes, 4 infectious diseases, and 1 cardiovascular phenotype. These results are consistent with higher comorbidity of diseases affecting inflammation, connective tissue, and the circulatory system in individuals with XFS **(Suppl. Table S10a, Suppl. Figure S6a)**.

XFS polygenic score was not significantly associated with any phenotypes in the analysis **(Suppl. Figure S6b)**. This potentially indicated that PRS generated from the global multi-ethnic GWAS summary might not be powered to detect association with traits in the EHR, and we might require scores from a more homogeneous and a much larger sample size. However, among the top PRS associations, we found several inflammatory diseases **(Suppl. Table S10b),** consistent with the enrichment results reported above.

## Discussion

We performed gene-based association analysis using GWAS summary statistics and conducted extensive experimental validation of genes associated with XFS. From our PrediXcan analysis, we identified 35 associated genes with XFS, 23 in single-tissue analysis and the rest in multi-tissue analysis. To eliminate the possibility of false-positive results due to LD contamination, we performed extensive additional analyses. First, we performed PrediXcan analysis in reduced models removing variants in LD with the two *LOXL1* missense variants associated with XFS, and variants in *LOXL1/STOML1* models in both global multiethnic and a subset of European ancestry individuals. Secondly, we conducted conditional analysis of the significant signals in European ancestry individuals. Thirdly, we then filtered signals based on correlated gene expression, LD and shared eQTLs and confirmed thirteen genes to be associated with XFS. Finally, expression analysis in human iris tissues further confirmed six of these seven signals, which were significantly downregulated in diseased XFS relative to normal eye tissues; *ARID3B, CD276, LOXL1, NEO1, SCAMP2 and UBL7*.

Our results suggest potentially substantial roles of inflammation and environment in the etiology of XFS. All of the six genes prioritized here by prediction and extensive validation analyses have inflammatory roles. *ARID3B, CD276, LOXL1* and *NEO1* are immunoregulatory molecules involved in the interaction between different tumors and the immune system.^59–62^ *SCAMP2* is important in granule exocytosis, a process crucial in membrane fusion in normal cellular functions in diverse systems including the immune system’s inflammatory response.^63–65^ *CD276* is involved in regulation of Ag-specific T cell-mediated immune responses and participates in the innate immunity-associated inflammatory response.^66,67^ *LOXL1* has also been implicated in fibrosis in response to inflammation in human breast cancer,^68^ in liver and lungs in model animals.^69–71^ *UBL7* encodes a member of the ubiquitin protein family, that is crucial in immune response and regulation of inflammatory response.^72–74^

Genes that show significant association of predicted expression with XFS at nominal significance are enriched for genes associated with three inflammatory conditions: rheumatoid arthritis, Type 1 diabetes and vitiligo in the Jensen Diseases database, with genes associated with the former two conditions enriched even with HLA region excluded. This is also consistent with the enrichment we find in DSigDB and DrugBank for cyclosporin A, acetaminophen and quercetin, which are compounds that have anti-inflammatory effects.^75^

Enrichment of predicted genes in this study in the polyunsaturated fatty acid (PUFA) and steroid derivatives: protectin (Reactome), omega fatty acid and estrogen (WikiPathways) are also consistent with the potential role of inflammation in XFS. Protectin, a derivative of PUFA including Omega-3 that are major components of fish oil, has an anti-inflammatory, anti- amyloidogenic, and anti-apoptotic activities in human neural cells.^76–78^ Omega fatty acid has been suggested as an IOP reducing supplements^79–81^ because of its anti-inflammatory effects.^82^ Association of steroid derivative, estrogen with glaucoma has been previously explored with higher levels of estrogen in reduction in IOP and conferring a possible reduced risk of glaucoma.^83^ The synthetic form of estrogen, estradiol, has been shown in a rat glaucoma model to inhibit optic nerve axonal degeneration by inducing a protein that is crucial in protecting RGC from oxidative damage.^84,85^

The association with inflammation is consistent with studies in limited numbers of XFS patients that found elevated inflammatory markers relative to controls, including cytokines, and markers such as interleukin-6 *(IL-6)* and *IL-8*,^86,87^ tumor necrosis factor-α *(TNF-α)* and *YKL- 40*.^88,89^ However, there are conflicting results for high sensitivity C-reactive protein.^90,91^

In addition, the XFS gene sets are enriched for genes that map to variants implicated in coffee and caffeine intake. Effects of caffeine consumption in the etiology of XFS have been studied, on the premise that coffee consumption increases plasma homocysteine levels that are speculated to enhance XFS material formation by contributing to vascular damage, oxidative stress, and extracellular matrix alterations.^92–95^ Consumption of coffee has been reported to have both pro- and anti-inflammatory effects.^96^ However, review of fifteen studies on the effect of coffee and caffeine on inflammation inferred the former had anti-inflammatory action, while the latter had complex effects on the inflammatory response with both proinflammatory and antiinflammatory responses reported.^97^ Caffeine might have a neuroprotective role by regulating pathways that produce inflammatory molecules via adenosine receptors in brain cells.^98,99^ Posttranscriptional regulation of LOXL1 gene expression has been also shown to be modulated by caffeine.^100^

Globally, our results of the six novel functionally validated genes also confirm the role of connective tissue involvement in the etiology of XFS. Aung, *et al*.,^13^ demonstrated the role of haplotypes that carry *LOXL1* XFS causal coding variants in upregulating extracellular matrix components such as elastin and fibrillin, and increasing cell-cell adhesion. In addition, two of the novel genes in our study, *ARID3B* and *NEO1,* among the other six genes identified and validated in both studies, have adhesive roles in the body. *ARID3B* in conjunction with *FDZ5* protein increases adhesion to ECM components, collagen IV, fibronectin and vitronectin, that are components of exfoliation deposits.^101,102^ *NEO1* has also been shown to play adhesive role during organogenesis.^103^

Results from our enrichment analysis of genes associated with XFS are also consistent with a role of dysregulation in connective tissue metabolism in the etiology of XFS.

Cylosporin_A regulate *lysyl oxidase* expression and collagen metabolism probably by inhibiting an isomerase involved in protein folding.^104–106^ Other anti-inflammatory compounds identified from our enrichment analysis in the current study, epigallocatechin gallate, valproic acid, quercetin, ketoconazole and acetaminophen have also been shown to suppress collagen and/or are anti-fibrotic in variety of tissues by yet to be elucidated mechanism.^107–111^ Moreover, coffee and caffeine inhibit collagen expression and deposition, and have anti-fibrotic effects by blocking expressions and/or by modulating effects of profibrotic factors.^112–115^

Our results that show enrichment in crosslinking of collagen fibrils, a crucial constituent of connective tissues, and endosomal-vacuolar Reactome pathways, in our associated genes further confirm the importance of connective tissues in the etiology of XFS. In addition, there may be anomalies in an endosomal-vacuolar pathway shown to be involved in the accumulation of other aberrant proteins, including: Aβ peptides,^116^ prion^117^, and Huntingtin^118^ in neurons, and implicated in neurodegeneration. Moreover, inflammation has also been suggested in migratory failure and subsequent deposition of aberrant proteinaceous materials in affected tissues in conjunction with other molecular actors.^87,119–124^

Finally, our comorbidity analysis in the BioVU EHR indicated XFS association with several chronic inflammatory dermatological, musculo-skeletal, respiratory and infectious conditions. Moreover, extracellular matrix dysregulation is also suggested by our PheWAS results indicating XFS comorbidity with Vitamin D deficiency. Vitamin D regulate collagen cross-linking in vitro by upregulating gene expression of specific lysyl hydrogenase and oxidase enzymes.^125^

### Limitations of the study

This study has two main limitations. First, despite the fact that GTEx data for the 48 tissues represent the most comprehensive eQTL data set of human tissues, it does not constitute a complete representation of all human tissues and may fail to identify the real causal genes in the unsampled ocular tissue. However, we have confirmed from an ocular tissue database that novel signals identified in this study are robustly expressed in XFS relevant eye tissues. Moreover, recent analysis shows that the majority of the human body tissues exhibit higher degrees of tissue similarities.^126^ In addition, it has been shown that most complex conditions, including XFS, might actually manifest in many diverse tissues in the body.^126^

Second, only a third of the signals identified in the larger data were robustly confirmed in a European dataset at genome-wide significance. This raised the possibility that most of the initial signals identified an artefact of local LD leakage or shared eQTLs with the sentinel *LOXL1/STOML1* signal. Using statistical validation with reduced models including no SNPs in LD with sentinel variants, we confirmed associations independent of *LOXL1* for at least ten genes including seven that were experimentally validated. In addition, results from a recent study are consistent with two other association gene signals confirmed using multi-tissue analysis of European dataset and PrediXcan of reduced models in multi-ethnic global data, *ISLR2* and *STRA6.^2^ ISLR2* and *STRA6* are both significantly downregulated in tissues of XFS patients together with other key components of the *STRA6* receptor-driven Retinoic acid (RA) signaling pathway, and that siRNA-mediated downregulation of RA signaling induces upregulation of *LOXL1* and XFS-associated matrix genes in XFS-relevant cell types.^25^ These data indicate that dysregulation of *STRA6* and impaired retinoid metabolism are involved in the pathophysiology of XFS syndrome. Retinoic acid, the active metabolite involved in the signaling pathway implicated by Berner *et al*^25^ in XFS through regulation of *ISLR2, STRA6* and *LOXL1*, has been shown to control critical checkpoints in inflammation and to promote an inflammatory environment.^127–129^

In summary, our analysis of predicted gene expression and extensive functional analysis in eye tissue prioritized six genes in association with XFS. Our results further confirmed the role of connective tissues and highlighted the importance of inflammation in the etiology of XFS. Thus, molecular elements that underlie the interaction of connective tissue biosynthesis and inflammatory pathways may play a central role in the etiology of XFS.

## ACKNOWLEDGMENTS

We are grateful to Maria Niarchou and Tyne Fleming for comments on earlier version of the manuscript.

JH was jointly supported through grant to MAB (T32 grant 5T32EY021453), ERG and NJC.

ERG is grateful to the President and Fellows of Clare Hall, University of Cambridge for the fellowship support. ERG is also supported by a NIH Genomic Innovator Award (R35HG010718). The content is solely the responsibility of the authors and does not necessarily represent the official views of the National Institutes of Health.

KMJ - Joseph Ellis Family and William Black Research Funds, NEI Core Grant 6P30EY08126 to Vanderbilt Vision Research Center, Unrestricted Departmental Grant from Research to Prevent Blindness, Inc., NY.

The European-Data-Set was support by the Interdisciplinary Center for Clinical Research (IZKF) at the University Hospital of the University of Erlangen-Nuremberg (project E23) to AR and USS and the Deutsche Forschungsgemeinschaft (SCHL 366/8-1) to USS.

JH, NJC and ERG jointly conceived the project. AR, US and FP managed patients ‘ data and tissues’ samples of the three European cohorts. US, DB and FP conducted functional biological experiments. FP contributed raw genotyping data for European populations. JH performed all the statistical analysis. JH drafted the manuscript with critical input from KJ, NJC, ERG, FP & CCK. The manuscript was approved by all authors. NJC was responsible for obtaining financial support for this study.

